# The Micromechanical Environment of the Impinged Achilles Tendon Insertion

**DOI:** 10.1101/2022.09.25.509413

**Authors:** Keshia E. Mora, Samuel J. Mlawer, Alayna E. Loiselle, Mark R. Buckley

**Author notes:** **Corresponding author**, Keshia E. Mora, 601 Elmwood Avenue, PO Box 88, Rochester, NY 14620.

## Abstract

Mechanical deformation applied to tendon at the tissue-scale is transferred to the microscale — including the extracellular matrix (ECM), the pericellular matrix (PCM), the cell and the nucleus — through a process known as strain transfer. Microscale strains, in turn, trigger biological activity that plays an important role in the maintenance of tendon phenotype and homeostasis. Although tendon predominantly experiences longitudinal tensile forces, transverse forces due to bony impingement have been implicated in both physiological (e.g., maintenance of the tendon insertion) and pathophysiological (e.g. insertional Achilles tendinopathy) processes. However, to our knowledge, prior studies have not characterized the micromechanical strain environment in the context of tendon impingement. Therefore, the objective of this study was to characterize the micromechanical strain environment in the impinged Achilles tendon insertion using a novel mouse hindlimb explant model in combination with finite element (FE) modeling. We hypothesized that impingement would generate large magnitudes of transverse compressive strain at the local matrix, PCM, and cell scales. Mouse hindlimb explants were imaged on a multiphoton microscope, and image stacks of the same population of tendon cells were obtained at the Achilles tendon insertion before and after dorsiflexion-induced impingement. Using an innovative multiphoton elastography approach, three-dimensional Green-Lagrange and principal strains were measured at the matrix scale, while longitudinal strain and aspect ratio were measured at the PCM and cell scales. Our results demonstrate that impingement generated substantial transverse compression at the matrix-scale, which led to longitudinal stretching of cells, an increase in cell aspect ratio, and — surprisingly — longitudinal *compression* of the tendon PCM. These experimental results were corroborated by an FE model developed to simulate the micromechanical environment in impinged regions of the Achilles tendon. Moreover, in both experiments and simulations, impingement-generated microscale stresses and strains were highly dependent on initial cell-cell gap spacing. Understanding the factors that influence the microscale strain environment generated by impingement could contribute to a more mechanistic understanding of impingement-induced tendinopathies and inform the development of approaches that disrupt the progression of pathology.

## Introduction

Maintenance of appropriate mechanical stimuli is crucial for preserving tendon homeostasis. For example, physiological tensile strains applied along the longitudinal axis of the tendon have been shown to preserve characteristics of healthy fibrous tendon including an aligned extracellular matrix (ECM), elongated cells, and high levels of type I collagen. In contrast, tendons subject to mechanical impingement — indentation due to contact with neighboring bony structures — adopt and maintain fibrocartilage-like features such as increased matrix disorganization, rounded chondrocyte-like cells, and elevated levels of type II collagen and aggrecan [1–3]. While biological adaptations to *physiological* impingement may be protective and may play a role in the formation of the tendon-to-bone attachment (enthesis), *excessive* impingement has been implicated in several tendon disorders including rotator cuff tendinopathy and insertional Achilles tendinopathy [4, 5]. However, the underlying mechanism explaining how mechanical impingement drives and preserves biological adaptations is poorly understood.

According to the paradigm of mechanobiology, tissue-scale strains are transferred to microscale strains at the matrix, pericellular matrix, cell and nuclear length scales through a process often described as “strain transfer”. Through several well-established mechanisms including ion channel activation [6, 7], chromatin condensation [8, 9], and integrin signaling [10, 11], these microscale strains, in turn, generate a biological response. Therefore, characterizing impingement-associated mechanical stimuli and determining how tissue-scale strains are translated into microscale strains is a critical step towards gaining a complete picture of how impingement triggers biological adaptations. However, due to the complex mechanical environment in impinged regions of tendon, tissue-scale strain patterns in these areas are often complex and difficult to predict. Moreover, previous studies in musculoskeletal tissue have demonstrated that the strain transfer response is highly heterogenous and tissue-specific, and tissue-scale strains transferred to the microscale can either be amplified or attenuated [12–14]. Notably, to our knowledge, strain transfer in the context of the multiaxial loading environment generated by impingement — an environment that includes both axial tension and transverse compressive strain — has not been previously explored in tendon.

Rigorously characterizing strain transfer in impinged tendon regions requires methods to accurately quantify the induced microscale strains in 3-D. Prior studies have implemented 2-D approaches to quantify microscale matrix strains including the use of photobleached lines and other external labels as fiduciary markers [15–17], as well as the tracking of in-plane nuclear centroids [18, 19]. However, these approaches lack the ability to capture the complexity of the microscale strain environment in its entirety, including out-of-plane deformations. Since the microscale strain environment dictates biological outcomes, having a more nuanced understanding of how microscale structures are perturbed by external loads ultimately aids in connecting mechanics to cell biology and tissue phenotype. A 3-D multiphoton elastography approach that enables simultaneous mapping of microscale strains at a high resolution across the local matrix, PCM and cell scales allows greater insight into how impingement uniquely affects microscale structures.

As a first step towards understanding how impingement drives the unique phenotype of impinged tendon regions such as the tendon-to-bone insertion, the overall objective of this study is to characterize the micromechanical strain environment in the impinged Achilles tendon. In prior ultrasound elastography studies [20], we demonstrated that at the tissue-scale, the dorsiflexed mouse Achilles tendon insertion experiences locally elevated magnitudes of transverse compressive strain, indicating impingement from the calcaneus — a finding that was consistent with previously studies in human subjects [21, 22]. An advantage of this explant dorsiflexion model is that it is highly controllable, yet — unlike other *in vitro* impingement models [23] — it preserves interactions between the Achilles tendon and surrounding tissues (i.e., bone, bursa, muscle) which allows for replication of the complex geometric characteristics of *in vivo* tendon impingement. Therefore, in the current study, we implemented this model — in combination with finite element (FE) modeling — to characterize the micromechanical strain environment of the impinged Achilles tendon insertion. We hypothesized that impingement would generate large magnitudes of transverse compressive stain at the local matrix, PCM and cell scales. By visualizing and quantifying microscale deformations in regions of tendon impingement for the first time, this study will begin to shed light on the key events that contribute to the development of pathologies triggered by impingement.

## Materials and Methods

### Experimental study design

Multiphoton elastography was used to characterize the micromechanical strain environment of the impinged Achilles tendon insertion in 8 hindlimb explants from 12-16 week old male C57BL/6J mice. Multiphoton image stacks of the same population of tendon cells were acquired at the Achilles tendon insertion before and after dorsiflexion-induced impingement (**Fig. 1**), and used to quantify local matrix strain, PCM deformation and cell deformation.

**Figure 1:**
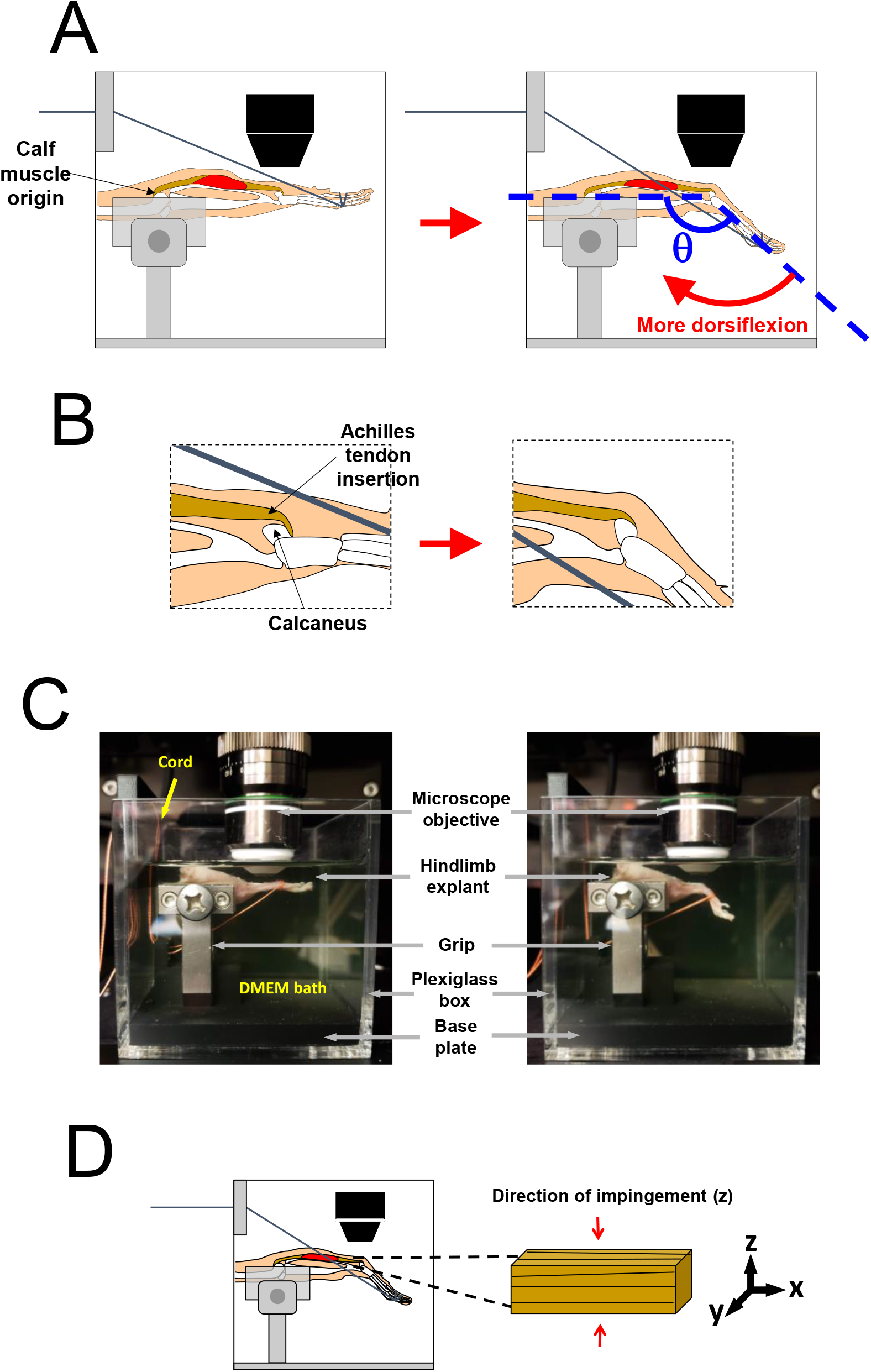
Experimental setup. **(A)** Schematic illustration of the multiphoton elastography platform before and after application of passive dorsiflexion by pulling a cord tied to the hindpaw of a mouse hindlimb explant. The ankle angle *θ* is the angle between the foot and tibia where the tibia and foot are parallel at an ankle angle of 180° and perpendicular at 90°. **(B)** Zoomed-in representation of the Achilles tendon insertion depicting how the calcaneus pushes against (i.e., “impinges” upon) the tendon after dorsiflexion. **(C)** Photograph of the multiphoton elastography platform with a representative mouse hindlimb explant at an initial 165° ankle angle (before impingement) and a final 125° ankle angle (after impingement). **(D)** Definition of the coordinate system used to describe directions within the Achilles tendon insertion. The *x*-axis lies along the fiber direction (the proximal-distal direction), the *y*-axis lies along the medial-lateral direction, and the *z*-axis lies along the direction of impingement (the anterior-posterior direction).

### Animal ethics

Experiments were carried out in accordance with the recommendations in the Guide for Care and Use of Laboratory Animals at the National Institutes of Health. All procedures performed on animals were completed at the University of Rochester and were approved by the University Committee on Animal Research (UCAR).

### Sample preparation

Hindlimb explants of 12-16 week old male C57BL/6J mice (N=8; The Jackson Laboratory, Bar Harbor, ME) were harvested at the femoral head with the overlying skin and adjacent plantaris tendon removed. Following dissection, explants were stained at 37°C on a rotator with fluorescein diacetate (FDA; 6 mg/mL, Thermo Fischer Scientific) for 1.5 hr, Hoechst 33342 (9.4 μM, Thermo Fischer Scientific) for 1.5 hr and propidium iodide (PI; 40 μg/mL, Thermo Fischer Scientific) for 15 min to visualize tenocyte cell intracellular space, to visualize nuclei, and to assess cell viability.

### Mechanical loading platform and specimen preparation

Fluorescently stained explants were loaded into a novel custom-built mechanical loading platform (**Fig. 1**). In this device, a sandpaper-coated stainless-steel grip (ADMET, Norwood, MA) was used to stabilize the mouse hindlimb explant with the knee in a fully extended position. This extended knee position introduced baseline longitudinal tension in the tendon, emulating contraction of the calf muscles. Next, the plexiglass container was filled with phenol red-free Dulbecco’s Modified Eagle Media (DMEM) that was preheated to 37°C to maintain hydration and viability of the explant throughout imaging. A cord tied around the foot was pulled to passively dorsiflex the mouse paw and induce controlled impingement of the Achilles tendon insertion to a target ankle angle of *θ* = 125°, where we define *θ* to be 180° when the tibia and foot are parallel and 90° when the foot and ankle are perpendicular (**Fig. 1a**).

### Multiphoton imaging

The Achilles tendon was imaged with a multiphoton microscope (Olympus FVMPE-RS) that interfaced with the loading platform. The imaging system includes two lasers (one InsightX3 laser and one MaiTai HP DeepSee Ti:Sapphire laser) tuned to 780 nm and 920 nm, respectively, and imaging was performed through a 25x objective lens. Prior to dorsiflexion, the surface of the calcaneus was identified under the microscope by its characteristic fibrocartilage appearance. The position of the calcaneus was registered, and the microscope stage was moved to 0.5 mm away along the longitudinal axis towards the calf muscle. This location was chosen because previous ultrasound elastography studies performed using this platform demonstrated that this insertional tendon region experiences elevated magnitudes of transverse compressive strain, indicating localized impingement of the tendon insertion [20]. An overview image stack was obtained to capture any unique features (i.e., distinct cell clusters, distinct cell shapes) that could subsequently be used to identify the region of interest (ROI). A 3x optical zoom was applied, and a pre-impingement image stack was obtained. The acquired image stacks were 512 x 512 pixels (169.7 x 169.7 μm) with a 1 μm vertical (perpendicular to the imaging plane) step size (**Fig. 2**). Next, the explant was passively dorsiflexed by pulling the cord attached to the hindpaw. A protractor placed against the plexiglass cube was used to ensure that the hindpaw was dorsiflexed to a target final ankle angle of 125°. The ROI was subsequently located in its new position, and a post-impingement image stack was obtained using the same imaging settings.

**Figure 2:**
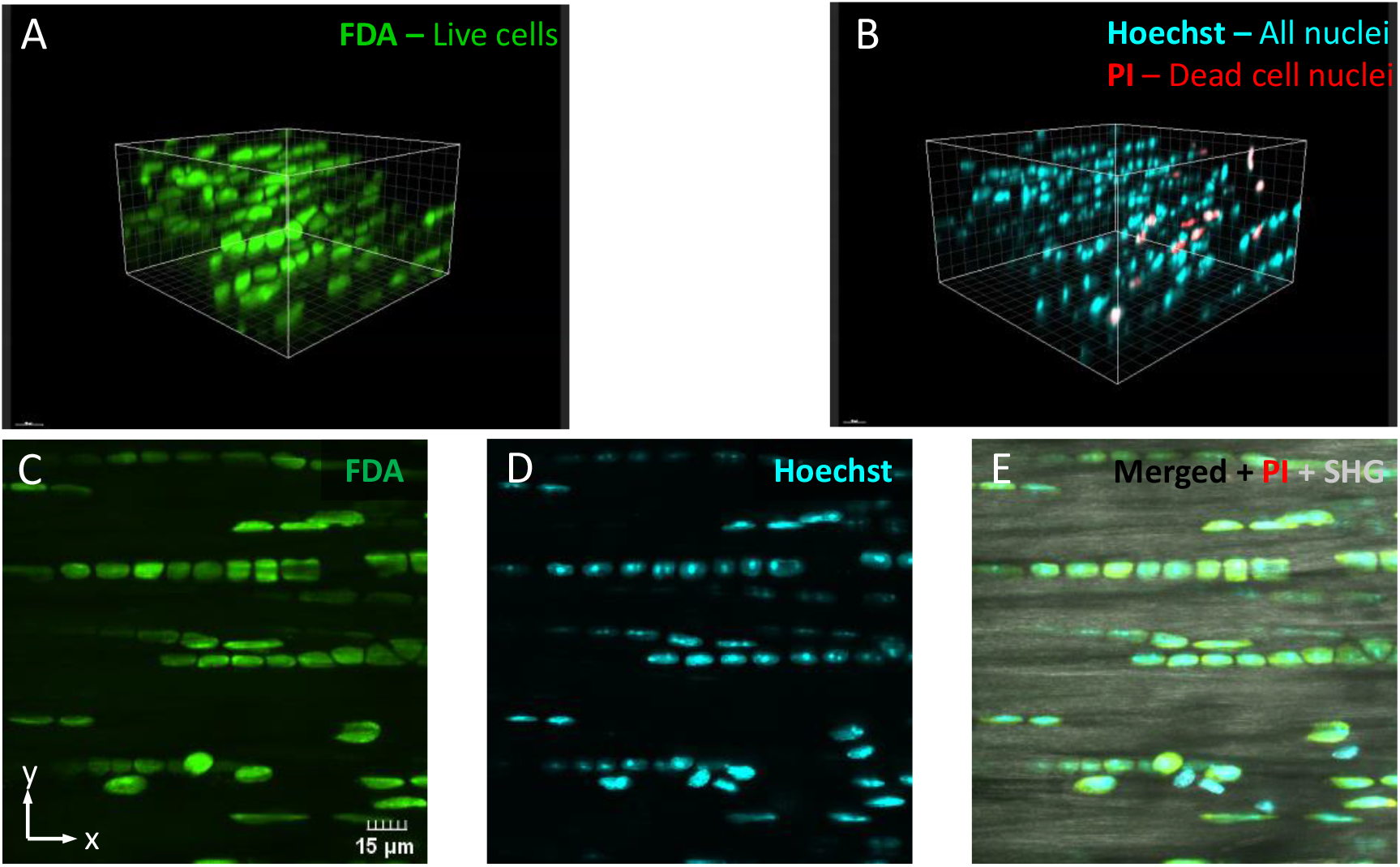
Representative multiphoton image stack. Representative multichannel multiphoton image stack of the mouse Achilles tendon insertion (0.5 mm from the calcaneus). **(A)** 3-D rendering of FDA-labelled live cells (green) and **(B)** Hoechst 33342-labeled nuclei (blue) and PI-labeled dead cells (red). **(C-E)** Z-projections of **(C)** FDA-labelled cells, **(D)** Hoeschst 33342-labeled nuclei, and **(E)** a merged overlay (FDA, Hoechst, PI and SHG). The scale bar is 15 μm.

### Three-dimension segmentation of multiphoton image stacks

Multiphoton image stacks of FDA-labelled cells were segmented in Amira (Thermo Fischer Scientific, v.2020.2, Waltham, MA) in order to determine cell centroid position and cell dimensions before and after passive dorsiflexion. First, a median filter was applied to the image stack. Next, the automated global thresholding tool in combination with the TopHat tool was used to threshold the image stack. Following thresholding, the Marker-Based Watershed algorithm was applied to segment and separate individual cells within the image stack (**Fig. S1A**). This procedure yielded a three-dimensional rendering of segmented cells throughout the image stack (**Fig. S1B**).

### Quantification of local matrix strain

Based on the identified centroids of individually segmented cells, cell position was tracked in both loading states (before and after impingement) to determine the local matrix strain using a custom MATLAB (The Mathworks, Inc v.9.6, Natick, MA) code. Local matrix strain was quantified by assessing the individual components of the Green-Lagrange strain tensor (*E*_11_, *E*_12_, *E*_13_, *E*_22_, *E*_23_ and *E*_33_) based on changes in the positions of the tracked cells in a coordinate system where the *x*-axis is parallel to the proximal-distal direction (i.e., along the fiber axis/long axis), the *y*-axis is parallel to the medial-lateral direction, and the *z*-axis is parallel to the anterior-posterior direction (i.e., along the direction of impingement) (**Fig. 1D**). First, every combination of two cells was used to define a set of position vectors in the before impingement loading state. Next, every combination of three vectors ***a***’, ***b**’* and ***c***’ in the pre-impingement loading state was used to determine a value of the deformation gradient tensor *F* defined by:

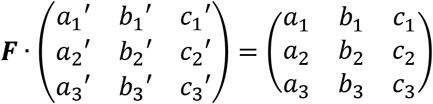

where ***a*** is the position of vector ***a***’ in the deformed configuration (i.e., after impingement). The peak of the distribution of each component of ***F*** (e.g., *F*_11_) was evaluated to determine a characteristic value of this component for the complete set of cells. Finally, the characteristic deformation gradient tensor ***F**_char_* — formed using the characteristic values of each component — was used to determine ***E*** according to:

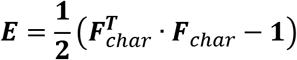

The maximum and minimum principal strains 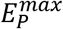 and 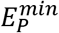 were evaluated by calculating the eigenvalues of ***E*** while the Jacobian ***I*** (indicating the relative volume change) was evaluated by calculating the determinant of ***F**_char_*.

### Quantification of pericellular matrix, cell, and nuclear deformation

The length of each segmented cell was taken to equal its maximum Feret’s diameter, i.e., the maximum distance between two parallel planes that are tangent to the cell. The minimum Feret’s diameter of each cell after the cell was projected onto a plane perpendicular to its length was taken to be the cell’s width. In general, the length axis was along or near the x-axis, while the width axis was along or near the y-axis. Cell strain along the length axis was computed as the change in cell length divided by the initial cell length. The cell aspect ratio (cell length divided by cell width; CAR) was also calculated. The cell-cell gap distance was taken to equal the minimum distance between adjacent cells within the same linear array, and the PCM strain was estimated as the change in cell-cell gap distance divided by the initial cell-cell gap distance.

### Finite element model geometry, mesh, and boundary conditions

To simulate the microscale strain environment generated by impingement, we constructed a quarter finite element (FE) model of a 75 x 75 x 15 μm region of the Achilles tendon insertion. The geometry was generated in Onshape and meshed using Gmsh, an open-source finite element mesh generator. Due to symmetry, only a quarter model was necessary to capture the 3-D behavior of the tendon region. The meshed geometry was then imported into the finite element modeling platform FEBio [24]. The microscale model contained separate parts representing the tendon extracellular matrix (ECM), pericellular matrix (PCM), and three linearly arranged cells within the PCM (**Fig. 3A**). The model mesh was biased, with a greater mesh density at and near the cell surface to capture local phenomena at the cell surface. In total, 398,504 tetrahedral elements and 75,463 nodes spanned the model, and this mesh size was determined based on a convergence analysis. Cell dimensions, PCM dimensions, and initial cell-cell spacing were set based on our acquired multiphoton image stacks as well as previous published electron microscopy studies [25]. Specifically, the cell length was 14 μm, the cell width was 7 μm, the PCM width was 9 μm, and the initial cell-cell gap distance varied between 0.5 and 7 μm (the approximate range of cell-cell gap distances in the mouse Achilles tendon insertion according to our multiphoton measurements). In the model coordinate system, the x-axis runs along the proximal-distal (longitudinal) direction, the y-axis runs along the medial-lateral direction, and the z-axis runs along the anterior-posterior direction (direction of impingement). To simulate impingement, a normal z-strain of −20%, a normal y-strain of −15% and a normal x-strain of 1% were imposed on the model (**Fig. 3B**). These values were consistent with the experimentally measured normal strains imposed by calcaneal impingement in our multiphoton elastography studies. For comparison, to simulate longitudinal tension, an x-strain of 10% was imposed on the model (**Fig. 3C**). Deformation along the y- and z-axes was unconstrained. Note that 10% strain is near the high end of the range of longitudinal strains experienced in the Achilles tendon insertion [22]. Finally, to simulate the combination of impingement and longitudinal tension, a normal z-strain of −20%, a normal y-strain of −15% and a normal x-strain of 10% were imposed on the model.

**Figure 3:**
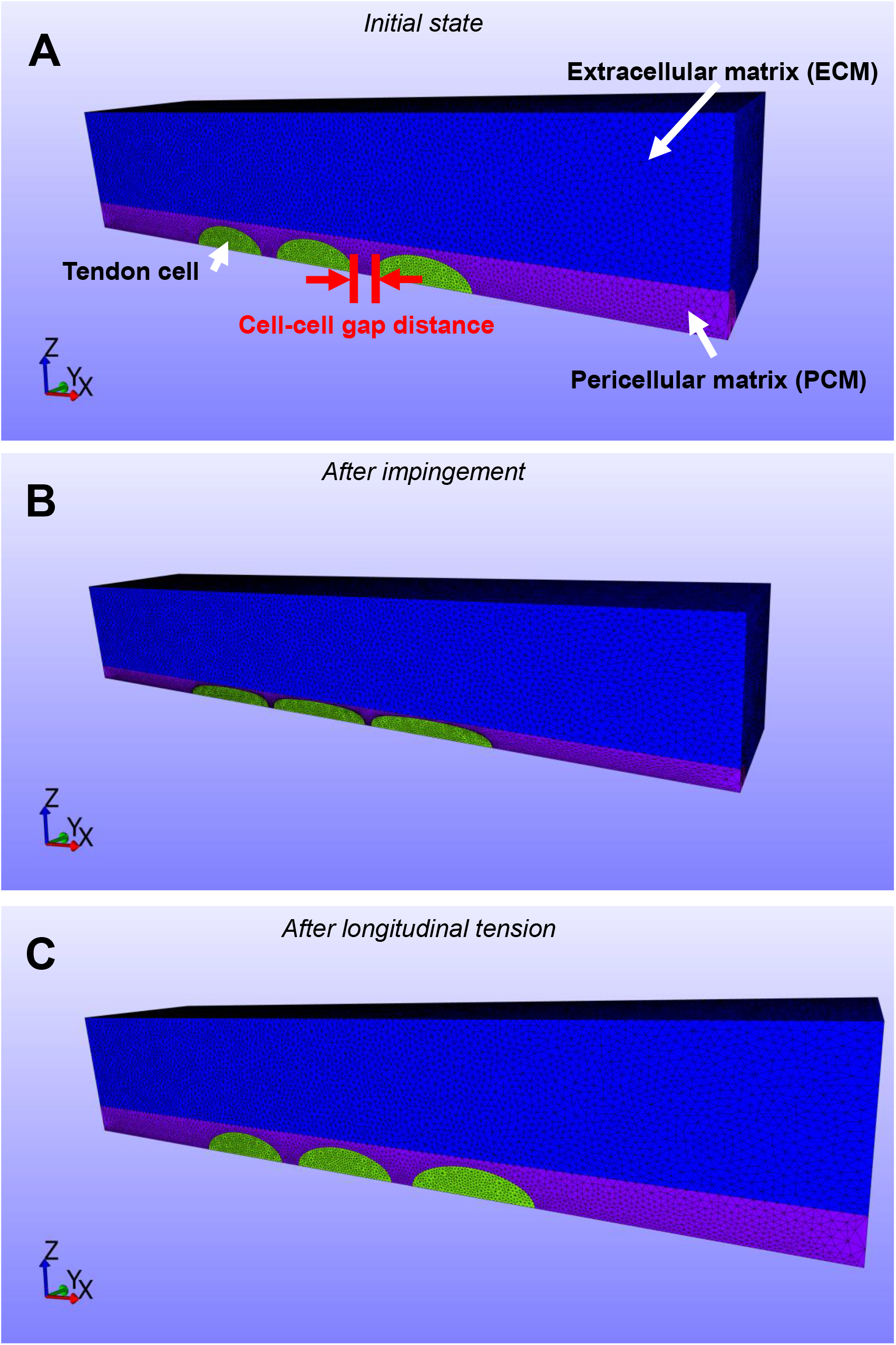
Microscale FE model of the impinged Achilles tendon insertion. Finite element (FE) model to stimulate the micromechanical strain environment in a small region (75 μm x 15 μm x 15 μm) of the impinged Achilles tendon insertion. **(A)** Meshed model geometry prior to loading including the ECM (blue), PCM (purple) and three cells (green). **(B)** Meshed model geometry after impingement (1% x-strain, −15% y-strain and −20% z-strain). Note the clear longitudinal expansion of cells along the x-axis and marked reduction in cell-cell gap distance induced by impingement. **(C)** Meshed model geometry after longitudinal tension (10% x-strain). Note the clear longitudinal expansion of cells along he x-axis without substantial reduction in cell-cell gap distance.

### Matrix constitutive model and material properties

The ECM was modeled as a transverse isotropic Mooney-Rivlin material [26–28] with fibers aligned along the longitudinal x-axis and material properties equivalent to the human medial collateral ligament [28]. This constitutive model uses a coupled strain energy function (i.e., a strain energy function with coupled deviatoric and dilatational components) given by:

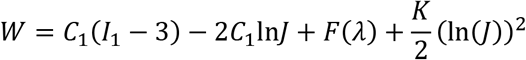

where the fiber response *F*(λ) satisfies:

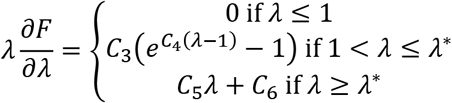

and

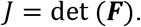

In the above equations, ***F*** is the deformation gradient tensor, ***I***_1_ is calculated from the right Cauchy green strain tensor ***C*** = ***F***^*T*^ · ***F*** according to:

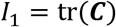

and the stretch ratio *λ* is calculated from ***C*** and the unit vector ***a***^0^ along the fiber direction according to:

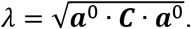

The second Piola-Kirchoff stress tensor *S* can be calculated from *W* using:

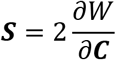

and from *S,* the true stress *σ* can be computed according to:

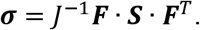

The 6 independent material properties of the transverse isotropic Mooney-Rivlin model are *C*_4_, *C*_3_, *C*_4_, *C*_5_, *K* and *λ*\*. *C*_4_ describes the mechanical response of the ground substance matrix, *C*_3_, *C*_4_,, *C*_5_ and *λ\** describe the mechanical response of the fibers, and *K* is the bulk modulus that describes the resistance of the material to volume change. Among the fiber properties, *λ\** is the stretch ratio at which the fibers become completely uncrimped (straightened), *C*_3_ is a scaling factor for the fiber stress (prior to uncrimping), *C*_4_ is the exponential growth constant for the fiber stress (prior to uncrimping), and *C*_5_ is the modulus of uncrimped fibers.

In a previous study, non-linear curve fitting of experimentally acquired stress-strain data was used to determine the parameters of the transverse isotropic Mooney Rivlin model for human medial collateral ligaments [28]. Specifically, the following values were measured: Λ* = 1.062, *C*_1_ = 1.44 *MPa, C*_3_ = 0.57 *MPa, C*_4_ = 48, and *C*_5_ = 467.1 *MPa.*

Due to the structural and compositional similarities of tendon and ligament, these parameter values were assigned to the ECM in our FE models. Furthermore, given the intent of the model to simulate equilibrium conditions (after transient load support from the pressurized interstitial fluid has ceased), the ECM was be assumed to be fully compressible and we took 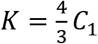.

### PCM constitutive model and material properties

The PCM was modeled as a (coupled) neo-Hookean material with properties equivalent to the PCM of articular cartilage. The constitutive equation for this model is:

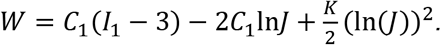

The material parameters *C*_1_ and *K* are related to the Young’s modulus *E* and the Poisson’s ratio *v* via:

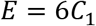

and

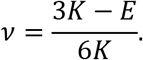

The mechanical properties of the PCM have not been assessed in tendon, but have been measured in articular cartilage — another collagenous musculoskeletal tissue — where it was found that *E* = 40 *kPa* and *v* = 0.04 [29]. In the current study, we assigned these values to the PCM in our FE model.

### Cell constitutive model and material properties

The cells were also modeled as neo-Hookean materials with properties equivalent to chondrocytes as measured in prior studies (*E* = 0.7 *kPa*) [29]. Since cells are nearly incompressible [30], the Poisson’s ratio of the cells was taken to be 0.499.

### Statistical analysis

Cell aspect ratio (CAR), cell length, cell width, and PCM length were each compared before and after impingement using a paired Student’s t-test with *α* = 0.05, while local matrix x-strain and cell x-strain were compared using an unpaired Student’s t-test with *α* = 0.05.

## Results

### Local matrix strain

Our data demonstrate that at the local matrix-scale, impingement generated relatively small (mean) longitudinal tensile strains of 1.2% long the x-axis (direction of collagen alignment; **Fig. 4A**). However, large transverse compressive strains (*E*_22_) of −10.5% were observed along the y-axis, likely due to the high Poisson’s ratio of tendon [31–33] (**Fig. 4B**). In addition, large transverse compressive strains (*E*_33_) of −20.0% were observed along the z-axis (direction of impingement; **Fig. 4C**). Small but variable shear strains were also generated at the local matrix-scale. Specifically, shear strains of 0.14% were observed along the y-z plane (*E*_23_; **Fig. 4D**), shear strains of −2.2% were observed along the x-z plane (*E*_13_; **Fig. 4E**), and shear strains of −1.7% were observed along the x-y plane (*E*_12_; **Fig. 4F**). Analysis of principal strains revealed a tensile maximum principal strain 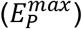 of 6.8% (**Fig. 4G**) and a compressive minimum principal strain 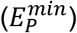 of −24.7% (**Fig. 4H**). Finally, the volume ratio, *J,* was 0.66, demonstrating that impingement resulted in marked fluid loss (**Fig. 4I**). Taken together these data demonstrate that our mouse impingement model generates large transverse compressive strains (*E*_22_, *E*_33_, 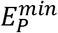) without substantial tensile strains (*E*_11_, 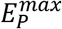) or shear strains (*E*_23_, *E*_13_, *E*_12_).

**Figure 4:**
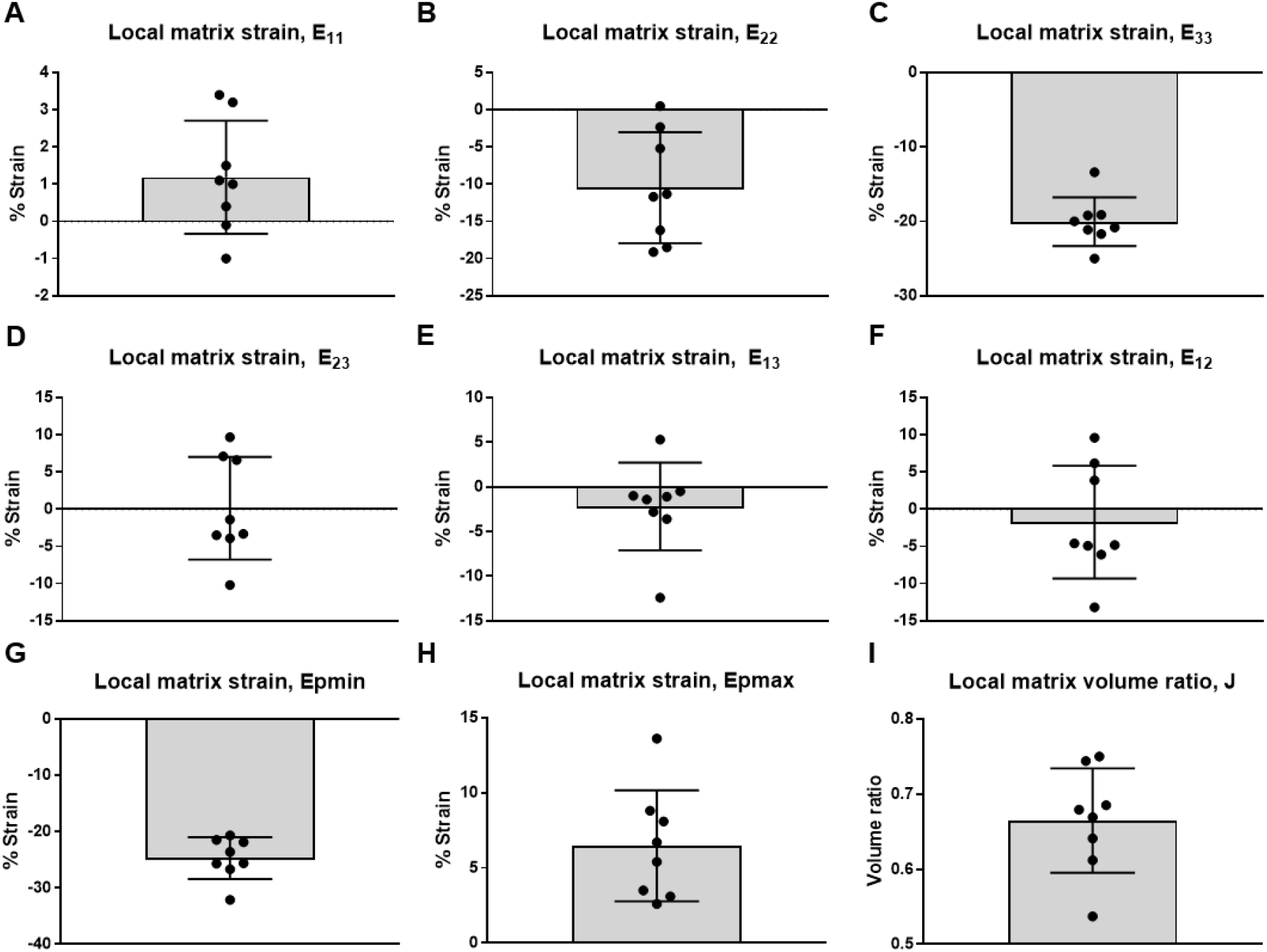
Local matrix strain generated by impingement as quantified through multiphoton elastography. Impingement-generated matrix strain within the insertion of the Achilles tendon (0.5 mm away from the postero-superior aspect of the calcaneus) was quantified using multiphoton elastography. Image stacks of the same population of tendon cells were obtained before and after impingement. Next, the Green-Lagrange strain tensor of the local matrix was calculated based on changes in tracked cell positions. Data demonstrate that impingement produced **(A)** relatively small (mean) longitudinal tensile strains of 1.2% along the x-axis (direction of collagen alignment), **(B)** transverse compressive strains of −10.5% along the y-axis (medial-lateral direction), and **(C)** large transverse compressive strains of −20.0% along the z-axis (direction of impingement). Impingement also generated relatively small shear strains of **(D)** 0.14% along the y-z plane, **(E)** −2.2% along the x-z plane, and **(F)** −1.7% along the x-y plane. Principal strains of **(G)** 6.8% and **(H)** −24.7% were induced by impingement, while **(I)** the relative volume change J was 0.66. Each data point represents the characteristic strain value for a single specimen based on analysis of an average of 24 cells/specimen.

### Pericellular matrix and cell deformation

At the cell-scale, the cell aspect ratio (cell length divided cell width; CAR) was increased after impingement (p=0.0051; **Fig. 5A-C**) due to both an increase in cell length (p=0.034; **Fig. 5D**) and a decrease in cell width (p=0.049; **Fig. 5E**). Due to cell elongation (increase in length along the longitudinal direction), the cell-cell gap between adjacent cells was reduced by impingement, resulting in approximately −27% compression of the PCM along the x-axis (**Fig. 5F**).

**Figure 5:**
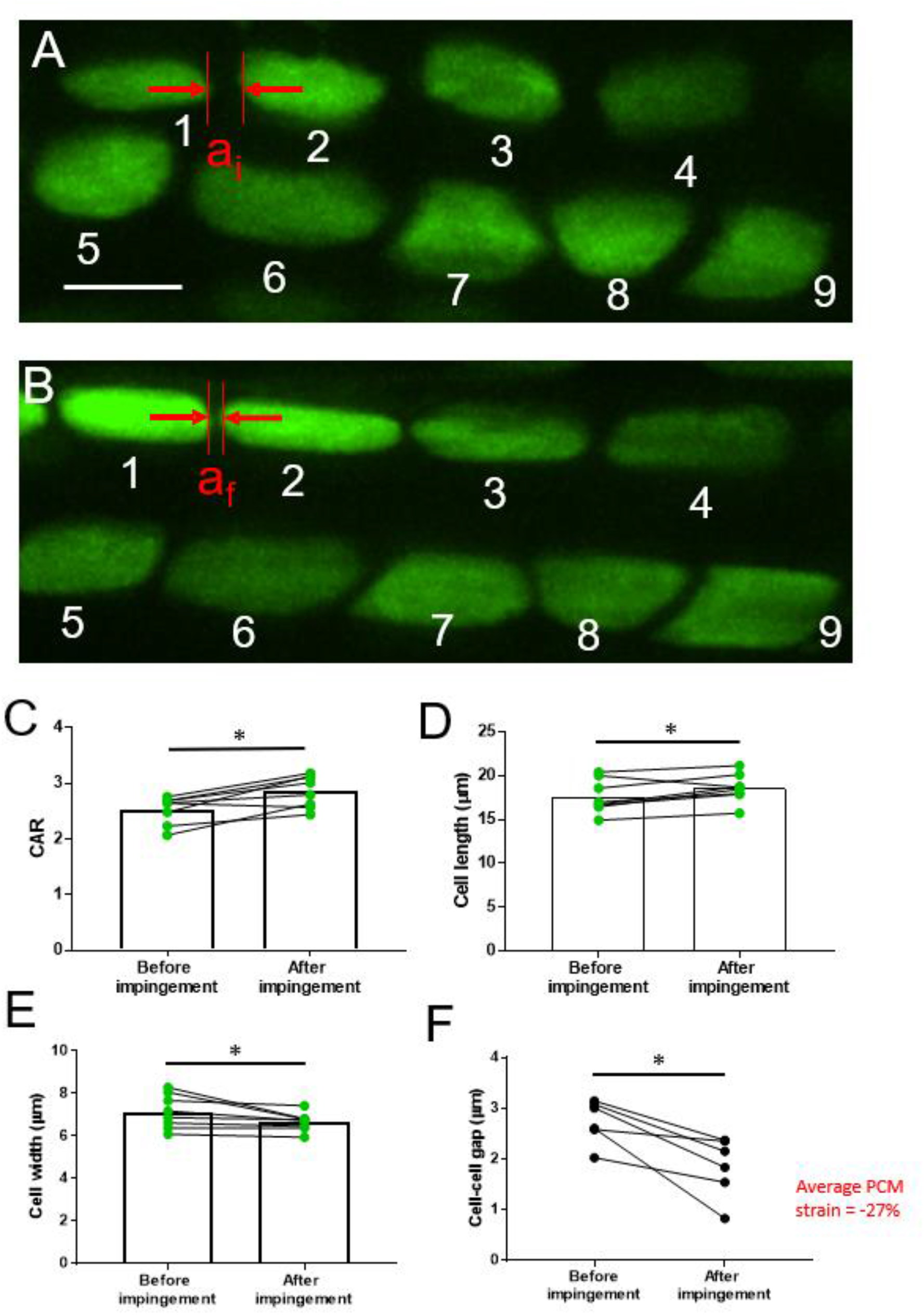
Cell and PCM deformation. **(A)** Representative z-projection of fluoresceine diacetate (FDA)-labelled cells before and after impingement, highlighting changes in cell aspect ratio, cell dimensions and cell-cell gap distance. In particular, note the marked reduction in the distance between cells 1 and 2 before («,) and after (*a_f_*) impingement. The scale bar is 10 μm. **(B)** Quantification of cell aspect ratio (CAR), demonstrating a significant increase in CAR after impingement. This increase in CAR is driven by **(C)** an increase in cell length and **(D)** a decrease in cell width. Each data point represents a single specimen and is calculated based on an analysis of an average of 24 cells/specimen. * indicates p<0.05. **(E)** Quantification of cell-cell gap distance demonstrates a significant reduction in cell-cell gap after impingement. Each data point represents a single specimen and is calculated based on analysis of an average of 8 cells/specimen. * indicates p<0.05. These data indicate that impingement produced an average longitudinal compressive strain of 27% in the tendon PCM.

### Strain amplification from the local matrix scale to the cell scale

At the local matrix scale, longitudinal tensile strain was low (1.2%; **Fig. 4A)**. However, at the cell scale, longitudinal strain (change in cell length divided by initial cell length) was 6.04%, significantly higher than at the local matrix scale (p =0.033; **Fig. 5D**). Hence, under impingement, we observed longitudinal strain amplification from the local matrix to the cell (**Fig. 6**). Importantly, this finding further corroborates the measured reduction in cell gap spacing and the measured longitudinal compression of the PCM, since longitudinal cell expansion without an equivalent elongation of the local matrix suggests a decrease in cell-cell gap distance and therefore compression of the PCM.

**Figure 6:**
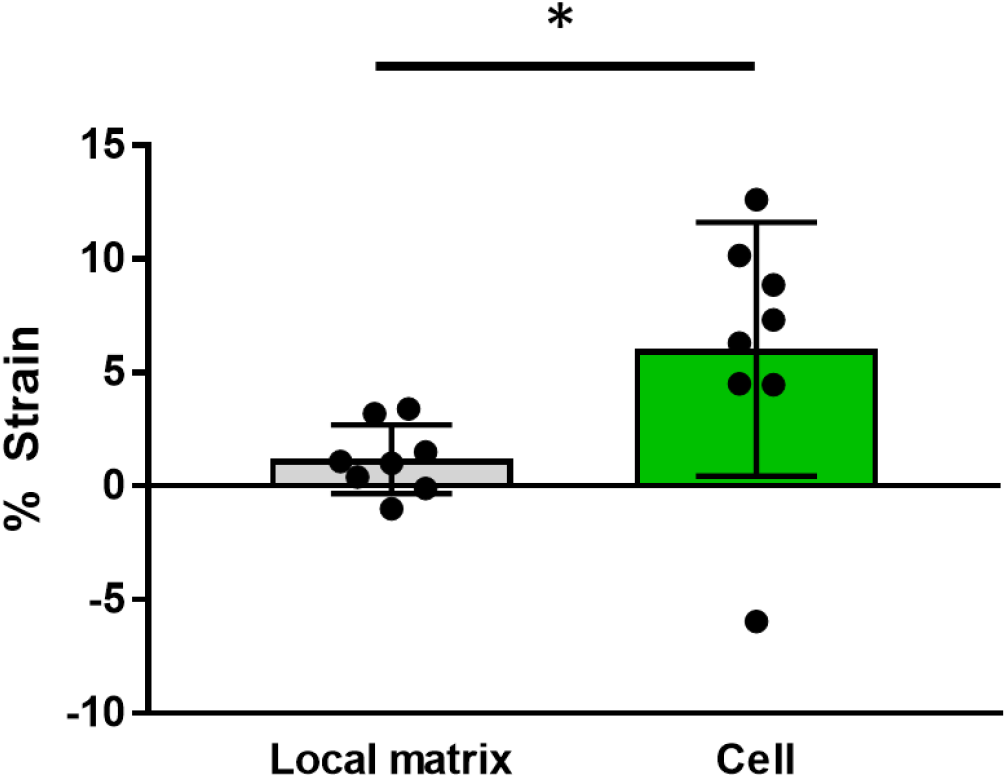
Local matrix and cell x-strain. Quantification of x-strains revealed longitudinal x-strain amplification from the local matrix scale (1.2%) to the cell scale (6.04%). * indicates p<0.05.

### Comparison with experimental studies

The finite element model reproduced several phenomena observed in our experimental multiphoton elastography studies. As in our experiments, a clear longitudinal expansion of cells (**Fig. 3B, Fig. 7A**) and a clear reduction in cell-cell gap distance (**Fig. 3B, Fig. 7B**) was observed after impingement. While the imposed x-strain on the ECM was just 1%, longitudinal cellular strains were greater than 1% for all initial cell-cell gap spacings and ranged between 7.5% and 23.3% (**Fig. 7A**), indicating substantial strain amplification from the ECM to cell scales as in experimental studies (**Fig. 6**).

**Figure 7:**
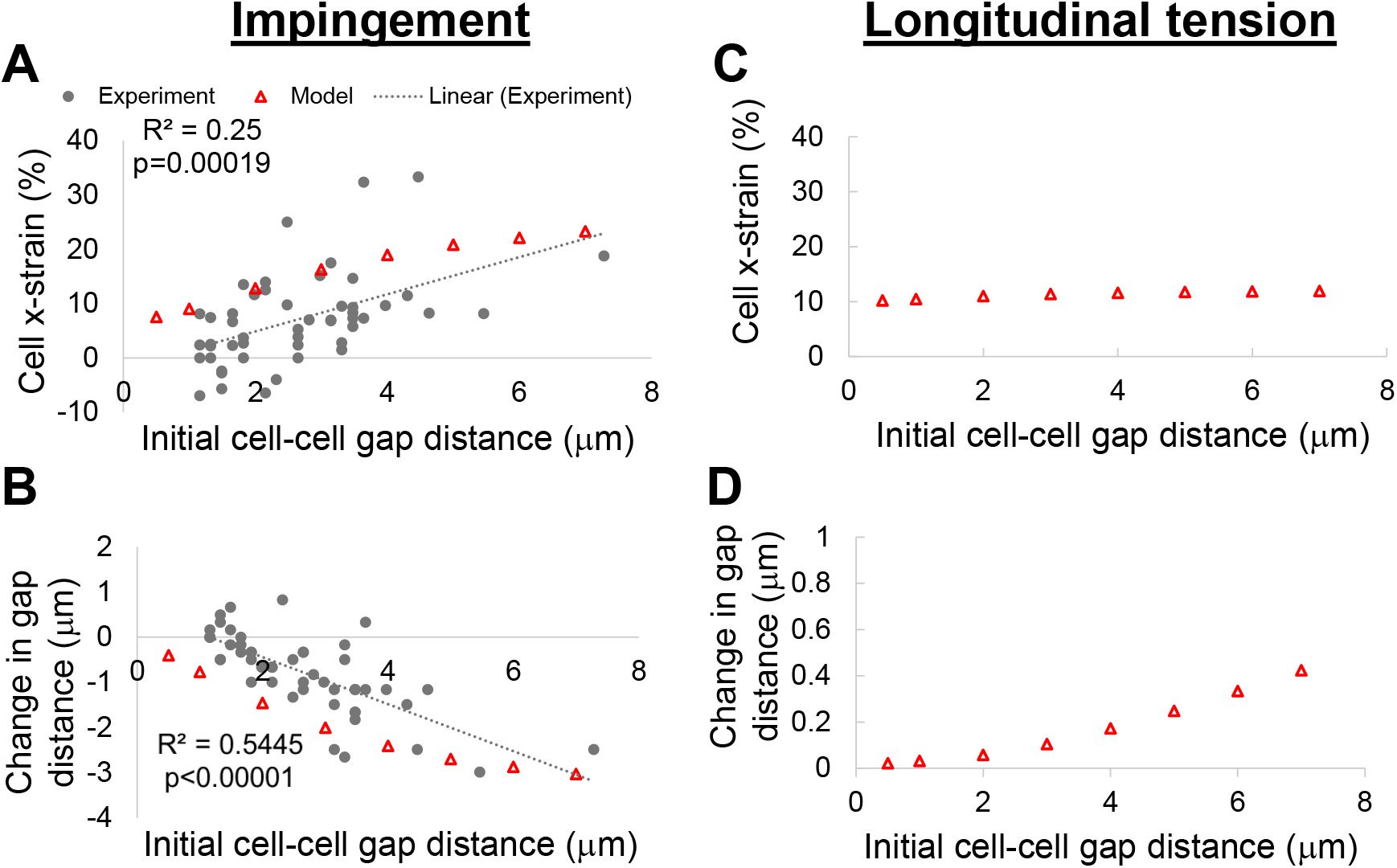
High cell x-strains and large reductions in cell-cell gap distance are generated by impingement and are correlated with initial cell-cell gap distance. **(A-B)** Relationships of initial cell-cell gap distance with **(A)** longitudinal cell strain and **(B)** change in cell-cell gap distance along the x-direction (fiber axis) after impingement (1% x- strain, −15% y-strain and −20% z-strain). Impingement induces high longitudinal cell strains that exceed ECM longitudinal strain (1%) and large reductions in cell-cell gap spacing (indicating PCM longitudinal compression). Moreover, according to both experimental data (gray circles) and the simulated FE model (red triangles), both cell x-strain and change in cell-cell gap distance are significantly correlated with initial cell-cell gap distance. **(C-D)** Relationships of initial cell-cell gap distance with **(C)** longitudinal cell strain and **(D)** change in cell-cell gap distance along the x-direction (fiber axis) after longitudinal tension (10% x-strain). Longitudinal tension induces cell x-strains that mirror ECM longitudinal strains (10%) for all initial cell-cell gap spacings. Unlike impingement, longitudinal tension causes a small increase (rather than a decrease) in cell-cell gap spacing.

According to the model, because cells with larger gap spacing have more room to deform, longitudinal cell strain (**Fig. 7A**) and reduction in cell-cell gap distance (**Fig. 7B**) after impingement both increased with increasing initial cell-cell gap distance. Remarkably, similar relationships were observed in our experiments (**Fig. 7A, B**). Hence, experimentally measured changes in microscale strain/deformation parameters with altered cell-cell spacing are captured by the FE model, both qualitatively and quantitatively.

### General comparison of impingement and longitudinal tension

Under impingement, the cells and the PCM experienced substantial compression along the z-direction (the direction of impingement) (**Fig. 3B**). But while impingement concomitantly caused the cells to expand longitudinally (along the x-direction) (**Fig. 3B**, **Fig. 7A**), the PCM separating adjacent cells was *compressed* longitudinally, leading to reduced cell-cell gap spacing (**Fig. 3B**, **Fig. 7B**). Under longitudinal tension, a different behavior was observed (**Fig. 3C**). The cells were stretched along the x-direction to a strain of approximately 10% (the same strain imposed on the model) (**Fig. 7C**). However, in contrast with impingement, the cell-cell spacing increased under longitudinal tension, reflecting stretching (rather than compression) of the PCM (**Fig. 7D**).

### Tensile strains at the cell surface are high under impingement

We applied our FE model to interrogate the strain environment in the impinged Achilles tendon at the sub-cellular scale, a scale that is challenging to assess experimentally. We found that cell surface tensile strain (i.e., 1st principle strain) was considerably higher for impingement compared with longitudinal tension and depended strongly on cell-cell gap spacing (**Fig. 8**). At low cell gap spacings, limited space for longitudinal cell expansion led to pronounced shape distortions and high local surface strains (**Fig. 8A**). However, when the gap distance was high, surface tensile strains were also high because each cell’s nearest neighbors are too far to inhibit cellular expansion (**Fig. 8C**). Thus, cell surface strain was minimized at a specific cell-cell spacing of approximately 3 μm (**Fig. 8B, F**).

**Figure 8:**
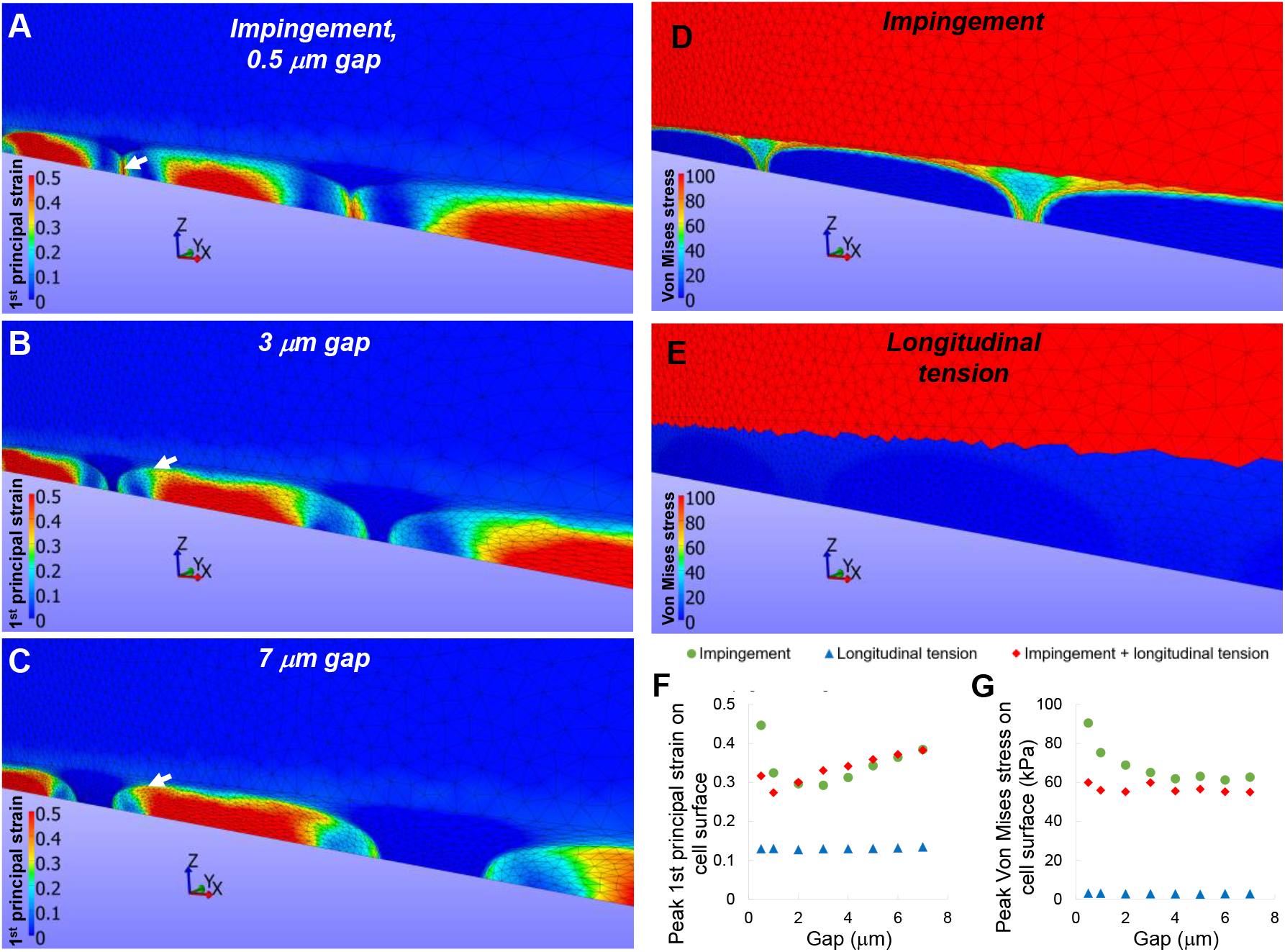
Distinct cell-scale stress and strain patterns are induced by impingement and longitudinal tension. **(A-C)** 1st principal strain maps for 0.5, 3 and 7 μm gap spacings. The location of the peak value along the surface of the middle cell is indicated by the white arrow. For small initial gaps (e.g., 0.5 μm), expanding cells are squeezed together and forced to undergo dramatic shape distortions that produce large strains on their proximal and distal edges. When the gap distance is larger (e.g., 3 μm), cells have more space to expand and surface strains are lower. But when the gap distance is very large (7 μm), cells have so much space to expand that high surface strains are again produced. **(D-E)** Von Mises stress maps for impingement and longitudinal tension (initial cell-cell gap = 5 μm). Under impingement, large stresses are generated at the cell surface. However, a similar phenomenon is not observed under longitudinal tension. **(F)** Quantification of peak 1st principal strain on the cell surface under impingement confirms high values at small and large initial cell gaps, leading to an optimal (minimum) value close to 3□m. In addition, surface strains are much lower for longitudinal tension only. **(G)** Quantification of Von mises stress on the cell surface demonstrates that higher surface stresses are generated by impingement when cell-cell gaps are small. In addition, surface stresses are dramatically lower for longitudinal tension compared with impingement.

### Von Mises stress at the cell border is high under impingement, but not under longitudinal tension

Motivated by the notable differences in cell surface strains for impingement as compared with longitudinal tension, we also used our model to assess cell surface stresses in the impinged and longitudinally stretched Achilles tendon insertion. Specifically, we evaluated and compared the spatially varying Von Mises stress in microscale regions of tendon subjected to impingement or longitudinal tension. The Von Mises stress is a scalar value of stress that reduces a complex, multiaxial stress state (e.g., the true stress tensor ***σ***) to a single value. It is often used to determine whether a material is likely to yield and is calculated according to:

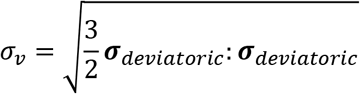

where ***σ**_deviatoric_* is the deviatoric part of *σ* given by

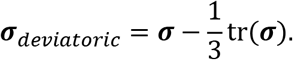

Our simulations demonstrated that under impingement, Von Mises stress in the cellular/pericellular region was highest at the cell surface. Moreover, the Von Mises stress at the cell surface under impingement was substantially greater than the Von Mises stress at the cell surface when longitudinal tension was simulated (**Fig. 8D, E, G**). Additionally, we found that the Von Mises stresses generated at the cell surface under impingement (but not longitudinal tension) were highly sensitive to initial cell-cell gap spacing. Specifically, a cell-cell gap distance less than 3 μm resulted in higher impingement-induced stresses at the cell border, as compared to cell-cell gap distances greater than 3 μm, which produced substantially lower impingement-induced stresses at cell border (**Fig. 8G**). Furthermore, for cell-cell gap distances greater than 3 μm the Von Mises stress at the cell surface induced by impingement appeared to plateau.

We also investigated how the combination of impingement and longitudinal tension — a strain state characterized by a normal z-strain of −20%, a normal y-strain of −15% and a normal x-strain of 10% — affected the cell surface Von Mises stress. We found that impingement and longitudinal tension together reduced cell surface stresses at all initial cell-cell gap distances compared with impingement only (**Fig. 8G**).

### Increased cell-cell gap distance as an adaptive response to impingement

Since our data indicate that cell-cell gap distance is an important mediator of cell-surface stresses and strains, we interrogated whether the tendon insertion, which is naturally impinged during ankle dorsiflexion, possessed altered cell-cell gap spacing compared to the midsubstance, which does not experience impingement. We found that the cell-cell gap distance at the insertion was approximately twice as large as the cell-cell gap distance at the midsubstance (**Fig. 9**). In addition, we also found that the mean cell-cell gap distance at the insertion was 3 μm, which is approximately equal to the cell-cell gap distance that minimizes cell surface tensile strain in our simulations (**Fig. 9**).

**Figure 9:**
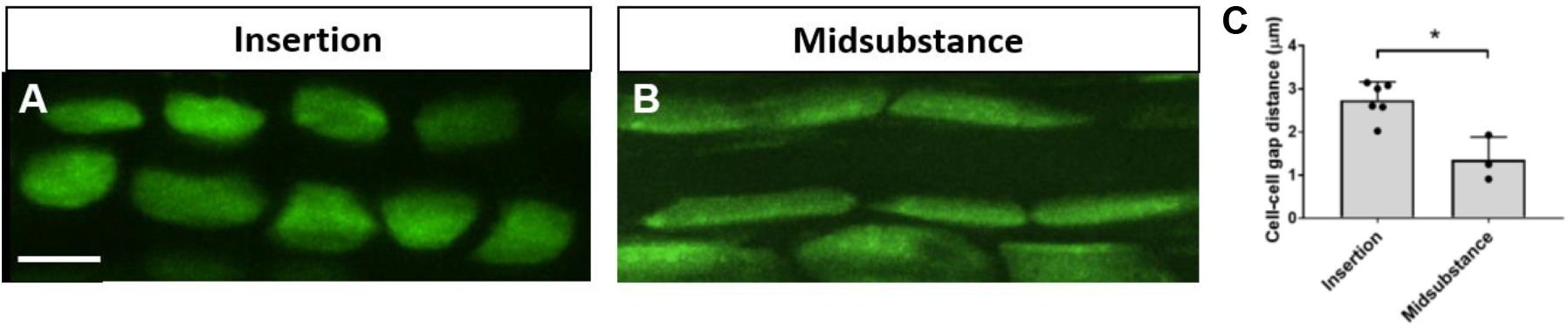
Increased cell-cell gap distance as an adaptive response to impingement. **(A-B)** Multiphoton micrographs of FDA-stained cells in the mouse Achilles tendon **(A)** insertion (0.5 mm from the calcaneus) and **(B)** midsubstance (2.5 mm from the calcaneus). The scale bar is 10 μm. **(C)** Cell-cell gaps are significantly larger in the insertion and are approximately equal to the value (3 μm) that minimizes cell surface tensile strain under impingement conditions.

## Discussion

To our knowledge, this is the first study to describe the micromechanical strain environment in the impinged tendon insertion. Specifically, we implemented a novel explant model and developed an innovative 3-D multiphoton elastography approach for high-resolution characterization of multi-scale strain patterns in the impinged Achilles tendon insertion. Our experimental findings demonstrated that impingement generates substantial transverse compressive strains at the local matrix scale, leading to cell elongation and longitudinal compression of the tendon PCM. Notably, both cell and PCM deformation were strongly dependent on cell-cell gap spacing, suggesting that cellular organization may in part govern the mechanobiological response to impingement. Importantly, a computational (FE) model corroborated all of our experimental findings and further demonstrated that impingement generates considerably higher cell surface stresses and strains as compared to longitudinal tension alone. However, our model suggested that combining longitudinal tension with impingement could mitigate high impingement-generated cell surface stresses.

At the local matrix scale, we demonstrated that impingement generates large transverse compressive strains without substantial longitudinal tensile strains or shear strains. These transverse compressive strains, in turn, squeeze water out of the tissue, resulting in a marked tissue volume loss of 34%. These findings are qualitatively consistent with the findings of our previous ultrasound elastography study [20]. However, in that study, slightly lower maximum principal strains (5% versus 6%) and lower transverse compressive strains (~12% versus ~25%) were reported (**Fig. S2**). These differences can be explained by the varying degrees of ankle dorsiflexion imposed in the two studies (dorsiflexion from 165° to 125° in the current study versus dorsiflexion from 145° to 125° in our ultrasound study [20]). In addition, a 2-D strain analysis was performed in our ultrasound elastography study while a 3-D strain analysis was performed in our multiphoton elastography study.

Both our experiments and simulations demonstrated that the length of the gap separating adjacent cells in a linear array is significantly decreased following impingement. Since the matrix in between tendon cells in linear arrays is contiguous and is part of the PCM [25], our data suggest that impingement longitudinally compresses the PCM in between tendon cells. This surprising microscale phenomenon is likely explained by the following: since tendon cells have a Poisson’s ratio near 0.5, they strongly resist volume changes and expand laterally when compressed. However, the PCM has a much lower Poisson’s ratio at equilibrium [29, 34], indicating that it tolerates volume changes and stretches less in lateral directions when compressed. Therefore, under impingement, compressed cells are forced to expand laterally and longitudinally compress the PCM. This finding may contribute to the mechanobiological response of tendon to impingement, by influencing cell-cell communication mediated within the PCM space.

Our modeling results also demonstrated that under impingement, tendon cell surface stresses and strains are substantially greater than under longitudinal tension. The distinct microscale stress/strain profile in impinged tendon regions could account for why *in vivo* impingement induces a unique biological response compared to longitudinal tension. Notably, when longitudinal tension was applied concomitantly with impingement, impingement-generated cell surface stresses were reduced, hinting that movements associated with high longitudinal tensile strains could mitigate the biological sequelae of chronic impingement. Presumably by increasing cell-cell gap spacing (**Fig. 7D**), longitudinal tension prevents cell-cell (mechanical) interactions that produce high cell surface stresses when cells in impinged regions are close to one another. Supporting this concept, we also found that impingement-induced cell surface stresses and strains are sensitive to initial cell-cell gap spacing, where small initial cell-cell gap spacing generated elevated cell surface stresses and strains. This phenomenon is likely explained by the following: cells with a very small initial cell-cell gap distance (e.g., 0.5 μm) have little space to expand laterally, forcing the cell to deform and distort into a shape that increases surface stresses and strains along the cell border. Therefore, cells that have a larger initial cell-cell gap distance (e.g., 3 μm), are shielded from high surface stresses and strains under impingement since they have space to laterally deform. However, if the initial cell-cell gap distance is too large (e.g., 7 μm), then high cell surface strains are generated because the cell has more space to stretch longitudinally since neighboring adjacent cells are too far away to inhibit cell expansion. Therefore, there appears to exist an optimal cell-cell gap distance of approximately 3 μm where cell surface strains are minimized.

Interestingly, we determined experimentally that the cell-cell gap spacing at the tendon insertion (which routinely experiences impingement) was approximately twice as large as the cell-cell gap spacing at the midsubstance (which generally remains unimpinged) (**Fig. 9**). We also found that the mean cell-cell gap distance at the insertion is 3 μm, which is approximately equal to the cell-cell gap distance that minimizes cell surface tension in our simulations (**Fig. 9**). Taken together, these observations imply that the increased cell-cell gap spacing of 3 μm observed at the tendon insertion may be the result of a protective adaptation (i.e., PCM growth) that allows cells in this region to withstand the stresses and strains induced by impingement.

Our 3-D multiphoton elastography approach is innovative because we simultaneous mapped 3-D strains across multiple length scales including the local matrix, PCM and cell scales at a high resolution and in a single experiment. In particular, to independently assess local matrix strain, we incorporated the position information of *all* tracked cells before and after impingement to define the deformation gradient tensor *F* and, in turn, the Lagrangian strain tensor *E* and the principal strains 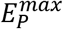 and 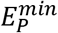. Our analysis has several key advantages compared to conventional 2-D approaches [15–17], or even certain 3-D approaches [35] used to assess microscale tissue strain patterns. Most of these studies applied digital volume correlation (DVC) [35] or other textural/correlation-based approaches to characterize strain patterns. However, these methods generally require large numbers of features (e.g., cells) to tabulate local displacements. Furthermore, the displacement field must then be translated into a strain field via spatial averaging, which is often susceptible to noise. In contrast, our method does not require spatial averaging and can work with either a small number of cells (4 cells is sufficient to compute a unique value of *F)* or large numbers of cells to allow averaging over broader regions.

A potential limitation of the experiments performed in this study is that the microscope is not able to penetrate the entire depth of Achilles tendon. Consequently, image stacks (between 50-60 μm in depth) were obtained at the tendon surface rather than in the part of the tendon closest to the calcaneus. For comparison, the total thickness of the tendon near the calcaneus is approximately 150 μm. However, based on previous ultrasound elastography studies where we detected elevated tissue-scale transverse compressive strains at the impinged tendon insertion *throughout the tendon depth* (due to the low thickness of the mouse Achilles tendon) [20], it is likely that microscale strains deeper in the tendon are similar to the microscale strains we measured at the tendon surface.

A limitation of the FE model in the current study is that the material properties that were incorporated into the model were based on published studies performed in ligament and cartilage, and not based on studies performed in mouse tendon fibrocartilage. To accurately assess cell, PCM and ECM material properties, future studies will implement atomic force microscope (AFM)-based nanoindentation testing of the mouse Achilles tendon insertion. The parameters obtained in these AFM studies will in turn be used to update our microscale FE model. Nevertheless, many of the experimentally observed behaviors at the microscale were recapitulated in the current model, which lends credibility to the current approach. Another potential limitation is that we have implemented a static model that considers the quasi-static response to mechanical deformation after viscoelastic and poroelastic effects in the cells, ECM and PCM have ceased. However, this approach allowed us to compare our model findings to our experimental multiphoton elastography studies, which evaluated microscale deformations in impinged tendon regions after equilibration (> 30 minutes after impingement). A final limitation of our FE model is that only normal micromechanical strains were imposed without accounting for shear. However, this approach is justified by the very small experimentally-measured shear strains in impinged regions of the Achilles tendon insertion (**Fig. 4D-F**).

In future studies, our multiphoton elastography approach can be used to understand how long-term impingement alters the microscale strain environment in either an *in vitro* explant model or an *in vivo* mouse model of Achilles tendon impingement. As the matrix closest to cells, the PCM is where matrix alterations first take place. For example, in cartilage, mechanical changes in the PCM are detectable just *3 days* after joint injury [36]. Therefore, the biological response to impingement could be detectable at early stages based on changes in the micromechanical strain environment of the PCM. Our multiphoton elastography can also be coupled with the use of transgenic mice and lineage tracing methods to understand if different tendon cell populations [37–39] undergo differential deformation under impingement. These studies could provide insight into cell populations that are more sensitive to impingement.

In conclusion, we characterized the microscale strain environment in an explant model of calcaneal impingement of the Achilles tendon insertion using an innovative 3-D multiphoton elastography approach. Both our experimental and modeling studies demonstrated that impingement generates distinct and complex patterns of microscale stress and strain that are absent in the context of longitudinal tensile loading alone. We also showed that high cell surface strains and stresses brought about by impingement are mitigated by modulating the arrangement of cells (e.g., through expansion of the PCM) or by applying longitudinal tension simultaneously with impingement. Since mechanical stimuli at the PCM and cellular scales are key steps in the chain of events leading from impingement to biological output, our findings shed light on the kinds of microscale changes that are likely to influence the biological sequelae of impingement. Therefore, this work could contribute to a more detailed mechanistic understanding of impingement-induced tendinopathies and ultimately inform the development of targeted approaches that disrupt strain transfer and inhibit the progression of pathology.

---

**Figure S1:**
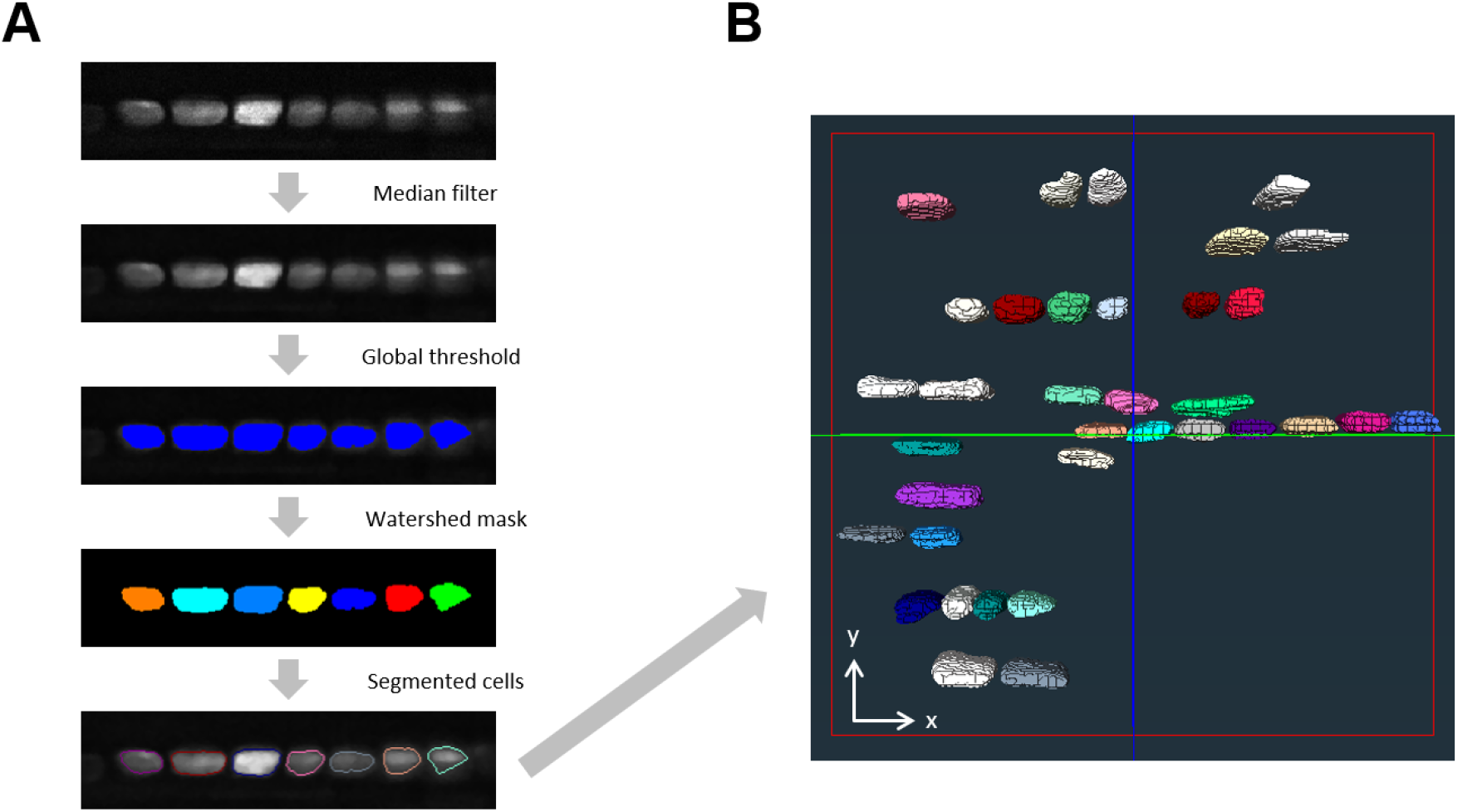
Three-dimension segmentation of multiphoton image stacks. Multiphoton image stacks of FDA-labelled cells were segmented in Amira to determine cell centroid position and cell dimension. **(A)** First, a median filter was applied to the image stack. Next, the automated global thresholding tool in combination with the TopHat tool was used to threshold the image stack. Following thresholding, the Marker-Based Watershed algorithm was applied to segment and separate individual cells within the image stack. **(B)** Rendering of tracked cells that were three-dimensionally segmented within a representative image stack.

**Figure S2:**
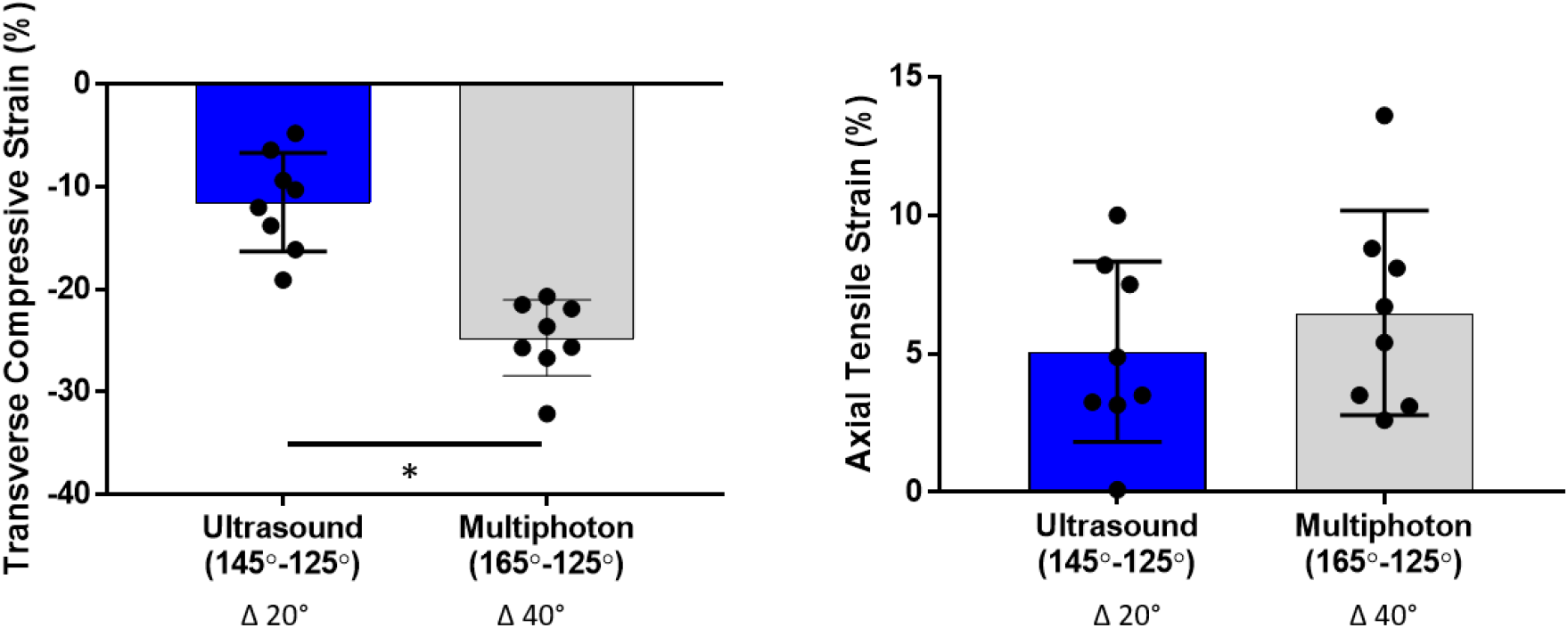
Comparison of ultrasound and multiphoton elastography-based measurements of matrix strain. Quantification of matrix strain was performed in both our ultrasound-elastography [20] and multiphoton elastography platforms. **(A)** Both studies were qualitatively consistent, demonstrating that the Achilles tendon insertion experiences marked transverse compressive strain (defined here as minimum principal strain) when the ankle is dorsiflexed. However, the magnitude of transverse compressive strain was higher in the current (multiphoton) study because the ankle was dorsiflexed to a greater extent (ankle angle change from 165° to 125° versus 145° to 125° in our ultrasound study). **(B)** Quantification of axial tensile strain (defined as maximum principal strain) was not significantly different between the two elastography-based measurements. * indicates p<0.0001.

